# Chronic kidney disease exacerbates hyperlipidemia and atherosclerosis via miR-148a-3p targeting SIK1/AMPKα1 in liver

**DOI:** 10.1101/2025.10.08.681287

**Authors:** Yuzhi Huang, Ting Li, Erwei Hu, Peiyao Zhang, Xueying Feng, Heze Fan, Zihao Wang, Yuyang Lei, Chen Wang, Wenbo Yang, Lijun Wang, Peining Liu, Bin Du, Yan Wu, Wenyuan Li, Jiahao Feng, Xiaofang Bai, Haoyu Wu, JuanLi Li, John Y-J. Shyy, Zuyi Yuan, Juan Zhou

## Abstract

**Background:** Cardiovascular complications are the leading cause of death in patients with chronic kidney disease (CKD). As a major risk factor for atherosclerotic cardiovascular disease (ASCVD), hyperlipidemia is a common sequela of CKD, which is linked to increased lipid synthesis, impaired lipoprotein clearance, and maladapted reverse cholesterol transport. Because exosomes can transport miRNAs to modulate intercellular and tissue communication, we hypothesized that exosomal miRNAs can act as endocrine-like molecules to modulate the CKD-associated hyperlipidemia and ASCVD.

**Methods:** Small RNA-seq and qPCR were used to determine the differential plasma level of exosomal miR-148a-3p in ASCVD patients with and without CKD. RNA-seq and *in silico* analysis were used to establish the regulation of lipid metabolism via miR-148a-3p/salt-inducible kinase 1 (SIK1)/AMP-activated protein kinase alpha 1 subunit (AMPKα1) in the liver. Inhibition of miR-148a-3p or overexpression of SIK1/AMPKα1 in ApoE^-/-^ mice with 5/6 nephrectomy was used to elucidate the role of the miR-148a-3p/SIK1/AMPKα1 axis in CKD-related hyperlipidemia and atherosclerosis.

**Results:** The plasma level of exosomal miR-148a-3p was significantly elevated in patients with ASCVD and CKD (ASCVD/CKD) and ApoE^-/-^ mice with 5/6 nephrectomy. Via circulating exosomes, kidney-generated miR-148a-3p was delivered to the liver, where it targeted *SIK1*/*AMPKα1* transcripts, thereby upregulating SREBP2/PCSK9 and reciprocally downregulating LDLR. Levels of SIK1 and AMPKα1 were decreased in hepatocytes transfected with miR-148a-3p or treated with plasma exosomes from ASCVD/CKD patients, with attendant SREBP2 activation and attenuated LDL-C binding. The decreased LDL-C binding was rectified by SIK1/AMPKα1 or LNA-miR-148a-3p overexpression. LNA-miR-148a-3p or AAV8-*SIK1*/*AMPKα* administration in 5/6 nephrectomy/ApoE^-/-^ mice significantly ameliorated hyperlipidemia and atherosclerosis.

**Conclusion:** In ASCVD/CKD patients and nephrectomy/ApoE^-/-^ mice, kidney-originating miR-148a-3p targeted hepatic *SIK1*/*AMPKα1* mRNA. This exosome-mediated endocrine effect upregulated SREBP2/PCSK9 and reciprocally reduced LDLR level, which elevated plasma LDL-C level and exacerbated ASCVD. These findings underscore a kidney-liver-artery axis involved in cardiovascular-kidney-metabolic syndrome.

## Introduction

Chronic kidney disease (CKD) has a broad impact on cardiovascular complications, including but not limited to atherosclerosis, myocardial infarction, and stroke.^1,2^ The American Heart Association defined this complex and interrelated systemic disorder as cardiovascular-kidney-metabolic (CKM) syndrome, which emphasizes interventions for metabolic syndrome-obesity, hyperlipidemia, insulin resistance and hypertension to reduce cardiovascular events associated with CKM.^3,4^

As a major risk factor of atherosclerotic cardiovascular disease (ASCVD), hyperlipidemia is a sequela of CKD.^5^ Among CKD patients, lipid disorders have a marked atherogenic profile, with elevated triglycerides (TG) and low-density lipoprotein-cholesterol (LDL-C) levels and reduced high-density lipoprotein-cholesterol (HDL-C) level.^6^ Thus, a subset of CKD patients are prescribed lipid-lowering drugs (e.g., statins and PCSK9 monoclonal antibody [mAb]) to maintain the lipid profile and mitigate cardiovascular incidents.^5,7^ Although lipid disorders are associated with clinical ramifications of CKD, the underlying mechanism remains elusive.

MicroRNAs (miRNAs) are small noncoding RNAs that regulate cellular function via a complementary interaction with the 3′-UTR of their targeted mRNA. Because of their vast involvement in physiology and diseases, miRNA drugs are emerging therapeutics that imitate or inhibit miRNAs.^8^ Relevant to CKM syndrome, miRNAs can be mediators linking CKD and CVD. For example, depletion of miR-16-5p, miR-17-5p, miR-20a-5p, and miR-106b-5p in CKD is associated with vascular calcification;^9^ miR-92a targets *SIRT1*, *KLF2*, and *KLF4* in vascular endothelial cells (ECs), which leads to endothelial dysfunction in CKD.^10^

MiRNAs can function as messengers between cells and tissues near and far.^11,12^ Exosomes, nano-sized extracellular vesicles, can transport miRNAs in the circulation for their endocrine-like action.^13,14^ However, we lack information on crucial miRNAs and their pathophysiologic role in CKM syndrome, particularly how nephrotic mRNAs affect hyperlipidemia and cause atherothrombotic diseases.

Here, we report a kidney-liver-artery axis involving miR-148a-3p that was known to induce atherosclerosis.^15-17^ The CKD-elevated miR-148a-3p is transported via exosomes to target hepatic *SIK1* and *AMPKα1* so that SREBP2 is desuppressed. With this endocrine-like axis, LDL-C level is elevated, which aggravates atherosclerosis. This novel mechanism may provide a rationale and therapeutic opportunity for CKD-related hyperlipidemia and ASCVD.

## Methods

### Animal studies

All animal experiments were conducted in accordance with the US National Institutes of Health (NIH) Guide for the Care and Use of Laboratory Animals and the Guidelines for Animal Experiments of Xi’an Jiaotong University Health Science Center. Animal experiments were approved by the Institutional Animal Ethics Committee of Xi’an Jiaotong University (XJTUAE2024-1892).

### Clinical studies

The clinical samples were obtained at the First Affiliated Hospital of Xi’an Jiaotong University with informed consent and institutional ethics committee approval (NO. 2021-1492). Detailed clinical characteristics of the ASCVD patients with or without CKD are provided in Table SI-III.

### Statistical analysis

Statistical analyses involved using GraphPad Prism 9 and data are expressed as mean±SEM. Initially, the datasets were analyzed for normality and equal variance to confirm the appropriateness of parametric tests. For data with normal distribution, 2-tailed Student t test (equal variance) or 2-tailed Student t test with Welch correction (unequal variance) was used to compare two groups and one-way ANOVA with Bonferroni post hoc test (equal variance) or Brown-Forsythe and Welch ANOVA test (unequal variance) to compare multiple groups. For non-normally distributed data, the Mann-Whitney U test was used to compare two groups and the Kruskal-Wallis test with Dunn post hoc test for multiple groups. Spearman’s rank correlation analysis was used to assess the correlation between levels of miR-148a-3p and other clinical biochemical indicators. The detailed statistical analysis for each experiment is presented in the corresponding figure legends. Biological experimental replicates in each group are shown in the figure legends. Differences with P < 0.05 were considered statistically significant.

### Data availability

All supporting data are available upon request.

Detailed description of the materials and methods is provided in the supplementary materials.

## Results

### Elevated miR-148a-3p Level in ASCVD Patients with CKD

To explore the pathophysiologic link between ASCVD and CKD via circulatory factors, we first profiled circulatory exosomal miRNAs in the discovery cohort containing two groups of patients: ASCVD with CKD (ASCVD/CKD, n=8) and matched ASCVD patients without CKD (n=8) (Figure 1A). The isolated exosomes, ranging from 30 to 200 nm, displayed typical features of exosomes (Figure S1). All exosome samples then underwent small RNA-seq. Overall, 51 miRNAs were differentially expressed between the two groups (Figure 1B). MiR-151-3p, -148a-3p, -423-5p, and -3184-3p were the most enriched among the 51 differentially expressed miRNAs (DEMs) (Figure 1C). On referring these 4 miRNAs to the Human microRNA Disease Database,^18^ only miR-148a-3p was associated with atherosclerosis in humans (Figure 1D). Because miR-148a-3p could regulate lipoprotein metabolism in mouse models,^19^ we recruited the second group of patients as a validation cohort to verify elevated miR-148a-3p level in human ASCVD/CKD. qPCR results confirmed that the plasma level of miR-148a-3p was inversely correlated with eGFR (i.e., stage of CKD) (Figure 1E). In addition, miR-148a-3p level was correlated with levels of TG, TC, and LDL-C in CKD patients (Figure 1F). Because urinary level of miRNAs can be used as a biomarker for kidney diseases,^20^ we measured the urinary level of miR-148-3p in the third ASCVD/CKD cohort. The urinary level of miR-148-3p was inversely correlated with eGFR (Figure 1G). Taken together, results in Figure 1 indicate that circulatory and urinary levels of miR-148a-3p were elevated in ASCVD/CKD patients.

**Figure 1.**
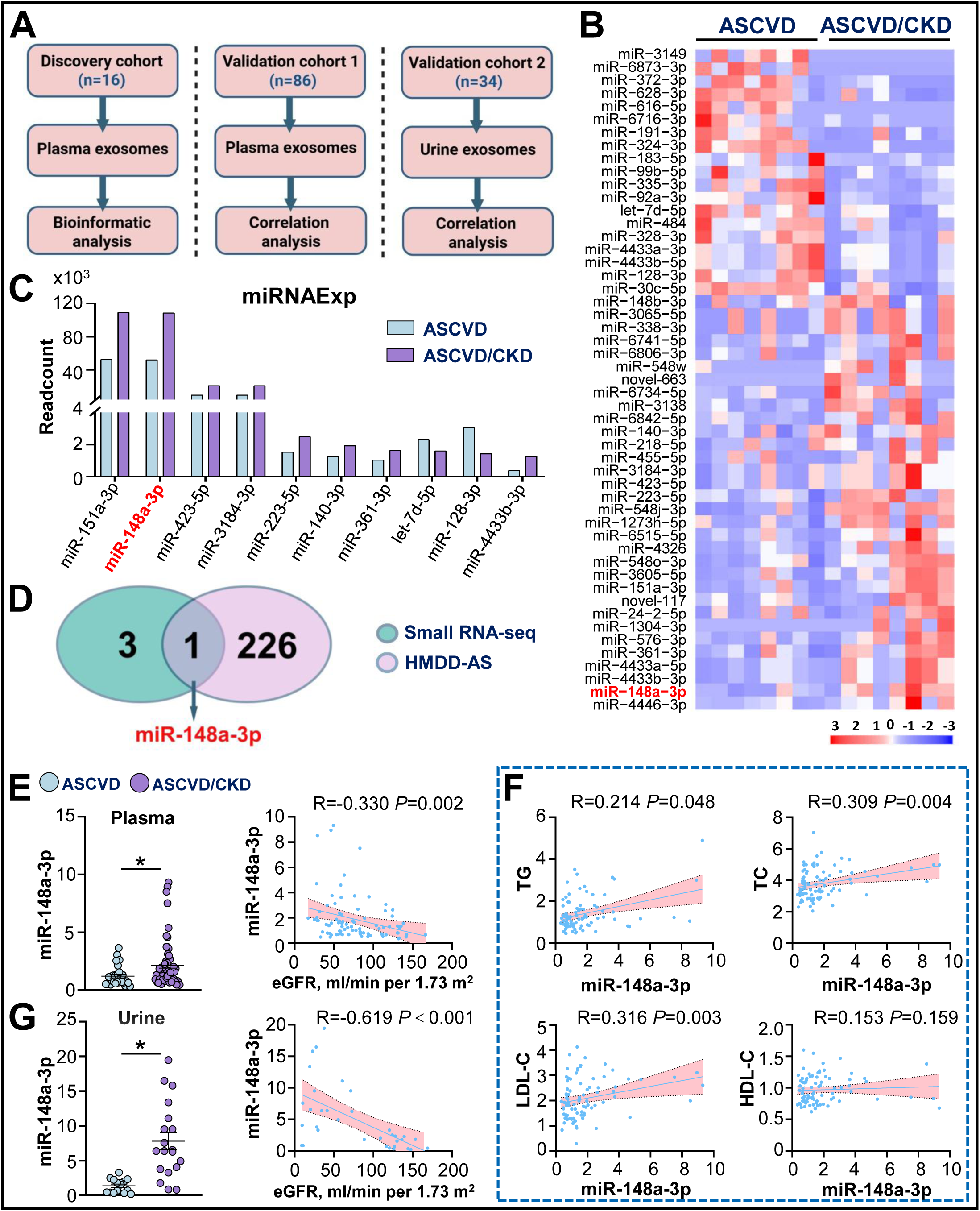
Elevated miR-148a-3p Level in Atherosclerotic Cardiovascular Disease (ASCVD) Patients with Chronic Kidney Disease (CKD). **(A**) In the discovery cohort, plasma exosomes from 8 ASCVD patients with CKD and 8 ASCVD patients without CKD were analyzed by small RNA-seq. The levels of plasma and urinary exosomal miRNAs were validated in validation cohort 1 comprising 28 ASCVD patients and 58 ASCVD/CKD patients and validation cohort 2 comprising 15 ASCVD patients and 19 ASCVD/CKD patients. **(B)** Heatmap of log10 (TPM+1) of the differentially expressed miRNAs (DEMs). **(C)** MiR-151a-3p, miR-148a-3p, miR-423-5p and miR-3184-3p were the most enriched among the top 10 DEMs. **(D)** Venn diagram demonstrating the common miRNAs in 2 databases: the top 4 enriched DEMs (green) and 227 atherosclerosis-associated miRNAs (HMDD v4.0, pink). **(E)** qPCR measurement of miR-148a-3p level in plasma exosomes from validation cohort 1. The data are fold-change normalized to the mean level in ASCVD patients. (**E, F)** Spearman analysis revealing correlations between miR-148a-3p level and eGFR as well as TG, TC, LDL-C, and HDL-C levels in validation cohort 1. **(G)** qPCR measurement of urinary exosome level of miR-148a-3p in validation cohort 2. The data are fold-change normalized to the mean level in ASCVD patients. Spearman analysis revealing correlation between miR-148a-3p level and eGFR in validation cohort 2. Normally distributed data were analyzed by 2-tailed Student *t* test with Welch correction (G) and non-normally distributed data by Mann-Whitney *U* test (E). \**P* < 0.05.

### LNA-miR-148a-3p Reduces Atherosclerosis in ApoE^-/-^/Nephrectomy Mice

To mimic human ASCVD/CKD in an animal model, we performed 5/6 nephrectomy in male ApoE^-/-^ mice (ApoE^-/-^/nephrectomy) fed a high-fat diet (HFD) (Figure 2A). In agreement with data from ASCVD/CKD patients, circulatory exosomal miR-148a-3p level was higher in hyperlipidemic ApoE^-/-^/nephrectomy mice than sham-operated ApoE^-/-^ mice fed an HFD (Figure 2B). Additionally, serum levels of Src, BUN, TG, TC, and LDL-C were elevated in male ApoE^-/-^/nephrectomy mice (Figure 2C, 2D). Corresponding to the elevated TC and LDL-C levels, atherosclerosis was enhanced in ApoE^-/-^/nephrectomy mice as compared with sham-operated controls (Figure 2E). To investigate the causality among nephrectomy-increased miR-148a-3p level and atherosclerosis, we administered LNA-miR-148a-3p to ApoE^-/-^/nephrectomy mice (Figure 2F). The level of miR-148a-3p in circulating exosomes, liver, and kidney was significantly lowered by LNA-miR-148a-3p treatment (Figure 2G and Figure S2A). With miR-148a-3p antagonism, levels of TG, TC, and LDL-C were lowered (Figure 2I). However, levels of BUN and Scr were not altered (Figure 2H). In accordance with the LNA-miR-148a-3p–decreased LDL-C level, atherosclerosis in ApoE^-/-^/nephrectomy mice was attenuated (Figure 2J). Furthermore, Oil-red O staining showed an increase in hepatic lipid content in these mice, which was ameliorated by LNA-miR-148a-3p treatment (Figure 2K). Of note, miR-148a-3p antagonism also reduced LDL-C level and atherosclerosis in female ApoE^-/-^/nephrectomy mice (Figure S2B-E). Taken together, data in Figure 2 suggest that miR-148a-3p could be a circulatory factor linking nephrectomy, hyperlipidemia, and atherosclerosis in mouse models mimicking human ASCVD/CKD.

**Figure 2.**
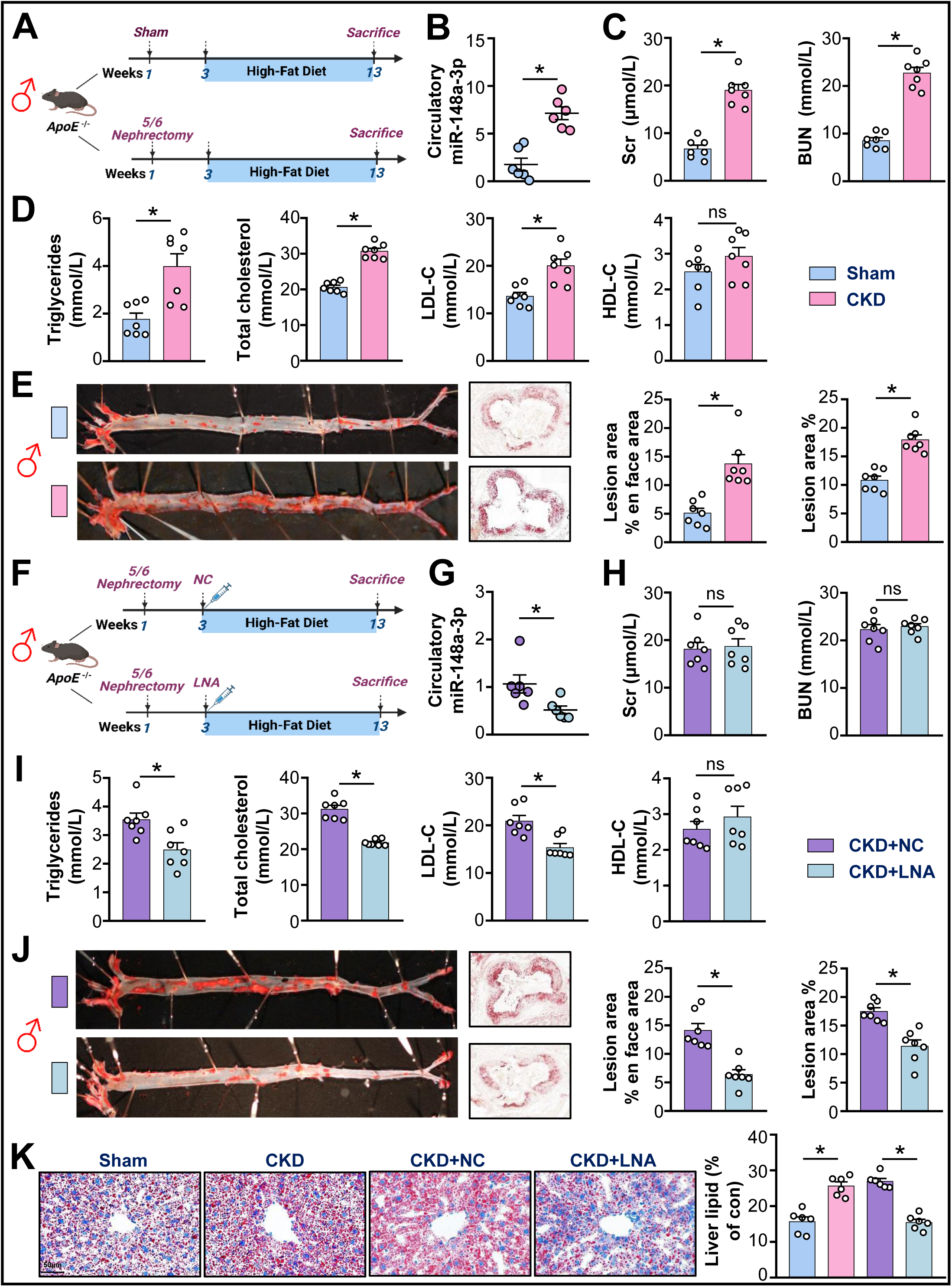
LNA-miR-148a-3p Mitigates Atherosclerosis in ApoE^-/-^/Nephrectomy Mice. **(A-E)** Eight-week-old male ApoE^-/-^ mice underwent 5/6 nephrectomy (CKD) or sham surgery (Sham), and were fed a high-fat diet (HFD) for 10 weeks. **(B)** miR-148a-3p level measured by qPCR in plasma exosomes from CKD and Sham mice. The data are fold-change normalized to the mean level in Sham controls. **(C, D)** Serum levels of Scr, BUN, TG, TC, LDL-C, and HDL-C in the two groups of mice. **(E)** Oil-red O *en face* staining of aorta and aortic root of ApoE^-/-^/nephrectomy and ApoE^-/-^/Sham mice. **(F-K)** ApoE^-/-^/nephrectomy mice fed an HFD received LNA-miR-148a-3p (CKD+LNA) or vehicle (CKD+NC) via tail-vein injection (20 mg/kg) bi-weekly for 10 weeks. **(G)** miR-148a-3p level in plasma exosomes from the two groups of mice. **(H, I)** Serum levels of Scr, BUN, TG, TC, LDL-C, and HDL-C. **(J)** Oil-red O *en face* staining of aorta and aortic root of ApoE^-/-^/nephrectomy mice with LNA or vehicle treatment. **(K)** Oil-red O staining of livers from Sham, CKD, CKD+NC, and CKD+LNA mice. Scale bar: 50 μm. Data are mean±SEM from 6-7 mice per group. Normally distributed data were analyzed by 2-tailed Student *t* test (B and C; TC, LDL-C and HDL-C in D; aortic root in E; H and J; TG in I) and non-normally distributed data by Mann-Whitney *U* test (TG in D; aorta in E; G; TC, LDL-C and HDL-C in I). Data were analyzed by one-way ANOVA among multiple groups in (K). \**P* < 0.05, ns=not significant.

### MiR-148a-3p Upregulates SREBP

Because of the striking changes in lipid profiles in ApoE^-/-^/nephrectomy mice, we administered LNA-miR-148a-3p to ApoE^-/-^/nephrectomy mice and used RNA-seq to study the transcriptomic variations in livers of these mice. LNA antagonism of miR-148a-3p upregulated 131 genes and downregulated 133 in mouse liver (Figure 3A). KEGG and GO analyses demonstrated that these differentially expressed genes (DEGs) are involved in pathways including metabolic pathways, bile secretion, steroid hormone biosynthesis, lipid and fatty acid metabolic process (Figure 3B and 3C). Given the relevance of these pathways to SREBP2-regulated cholesterol and bile acid metabolism as well as SREBP1c-regulated lipogenesis, we examined whether the expression of SREBP2, SREBP1c, and their regulated genes is negated by LNA-miR-148a-3p treatment. The mRNA level of *Srebp2* and its regulated HMG-CoA synthase, HMG-CoA reductase, and *Pcsk9*, together with *Srebp1c* and its regulated fatty acid synthase and acetyl carboxylase 1, were suppressed by LNA-miR-148a-3p (Figure 3D). Thus, miR-148a-3p might elevate LDL-C metabolism and hepatic lipogenesis via positive regulation of SREBP1,2. In support of this, LNA-miR-148a-3p administration to ApoE^-/-^/nephrectomy mice decreased the hepatic levels of mature-Srebp1,2; Pcsk9 (an SREBP2-regulated gene); and acetyl carboxylase 1 (an SREBP1c-regulated gene) but increased that of Ldlr (a PCSK9-downregulated gene) (Figure 3E, 3F). LNA-miR-148a-3p administration in these mice also reduced the circulatory level of PCSK9 (Figure 3G, 3H).

**Figure 3.**
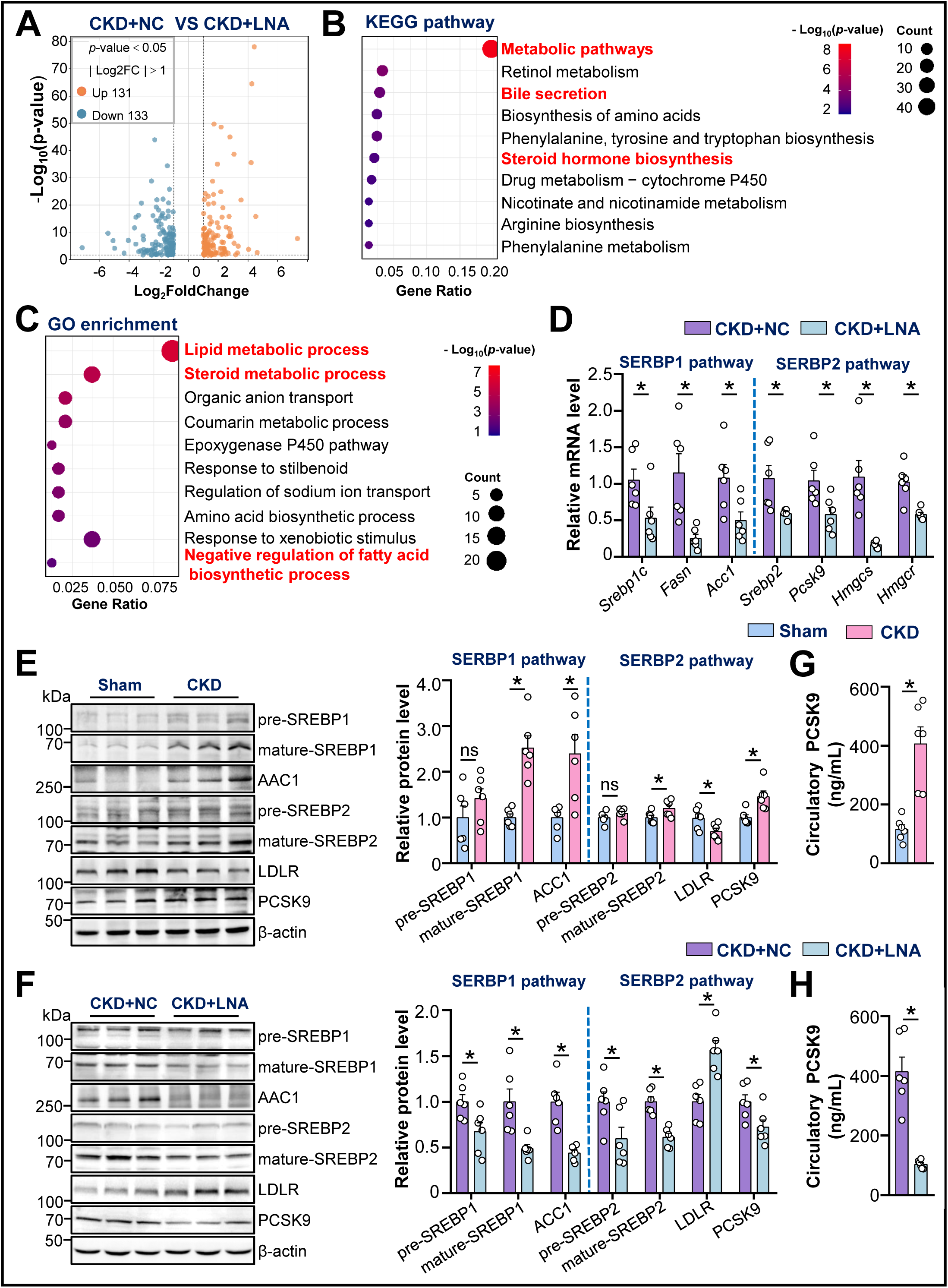
MiR-148a-3p Activation of SREBP in Hepatocytes. **(A)** Hepatic RNAs from ApoE^-/-^/nephrectomy mice administered LNA-miR-148a-3p or vehicle were analyzed by bulk RNA-seq. Volcano plot revealing DEGs in the two mouse groups (fold-change >1 or <-1, *P*<0.05). **(B, C)** KEGG enrichment pathway and Gene Ontology (GO) enrichment analyses using DAVID for the DEGs from RNA-seq, with cutoff log2 fold-change > 1 or < -1, *P* < 0.05. **(D)** mRNA levels of *Srebp1c*, *Srebp2*, and their regulated genes (*Fasn*, *Acc1*, *Pcsk9*, *Hmgcs*, and *Hmgcr*) in livers of ApoE^-/-^/nephrectomy mice treated with LNA-miR-148a-3p or vehicle. **(E, F)** Liver lysates from ApoE^-/-^/nephrectomy mice, sham-operated controls, and ApoE^-/-^/nephrectomy mice administered LNA-miR-148a-3p or vehicle were assayed by western blot analysis for SREBP1, ACC1, SREBP2, LDLR and PCSK9 levels, with β-actin as loading controls. **(G, H)** Plasma level of PCSK9 in the 4 groups of mice measured by ELISA. Data are mean±SEM from 6 independent experiments. Normally distributed data were analyzed by 2-tailed Student *t* test (Acc1 and Pcsk9 in D: pre-SREBP1, pre-SREBP2, mature-SREBP2, LDLR, PCSK9 in E and F) or 2-tailed Student t test with Welch correction (Fasn, Srebp2, Hmgcs, Hmgcr in D; mature-SREBP1 and ACC1 in E and F; G and H). Non-normally distributed data (Srebp1c in D) were analyzed by Mann-Whitney *U* test. \**P* < 0.05, ns=not significant.

### MiR-148a-3p Upregulation of SREBP2 via Targeting SIK1 and AMPKα1

Theorizing that miR-148a-3p upregulates SREBP2 and SREBP1 at the post-transcriptional level (i.e., miRNA targeting), we used Targetscan and miRDB to predict transcripts targeted by miR-148a-3p, which resulted in upregulated SREBP. The predicted targets were then referenced to hepatic transcriptomes obtained from ApoE^-/-^/nephrectomy mice (Figure 4A). This *in silico* analysis demonstrated that *Sik1, Gpcpd1, Esr1, Tfrc and Lrp4* were commonly suppressed by miR-148a-3p but restored by LNA-miR-148a-3p treatment. SIK1 belongs to the AMPK subfamily and is known to regulate glucose and lipid metabolism in liver.^21^ Although our RNA-seq data showed little effect of LNA-miR-148a-3p on AMPK, Targetscan analysis demonstrated a conserved miR-148a-3p-binding site in the 3′-UTR of *PRKAA1* (*AMPKα1*). The miR-148a-3p sequence and complementary seed sequences in the 3′-UTR of *SIK1* and *AMPKα1* are highly conserved among mammalian species, including human, rat, and mouse (Figure S3B). To this end, we constructed luciferase reporters encompassing luciferase fused to the 3′-UTR of *SIK1* or *AMPKα1* (*SIK1* 3′-UTR; *AMPKα1* 3′-UTR). As compared with control miRNA, miR-148a-3p overexpression in HEK293 cells decreased the luciferase activity of the co-transfected *SIK1* 3′-UTR (WT) or *AMPKα1* 3′-UTR (WT) (Figure 4B, 4C). However, miR-148a-3p–decreased luciferase activity was not seen in cells transfected with mutants of *SIK1* 3′-UTR and *AMPKα1* 3′-UTR (*SIK1* 3′-UTR [MUT]. *AMPKα1* 3 ′ -UTR [MUT]). To validate that miR-148a-3p can target *SIK1* and *AMPKα1* in hepatocytes, we treated HepG2 cells with miR-148a-3p, which decreased SIK1 and AMPKα1 protein and mRNA levels (Figure 4D). Additionally, immunoprecipitation experiments with anti-Argonaute-1 or anti-Argonaute-2 followed by PCR showed that miR-148a-3p overexpression in HepG2 cells increased *SIK1* and *AMPKα1* mRNA levels in the miRNA-induced silencing complexes (miRISCs) (Figure 4E). Function-wise, the hepatic levels of Sik1 and Ampkα1 were decreased in ApoE^-/-^/nephrectomy mice, which was reversed by LNA-miR-148a-3p administration (Figure 4F, 4G). Collectively, results in Figure 4 suggest that CKD-induced miR-148a-3p targeted hepatic *SIK1* and *AMPKα1*, which in turn upregulated SREBP.

**Figure 4.**
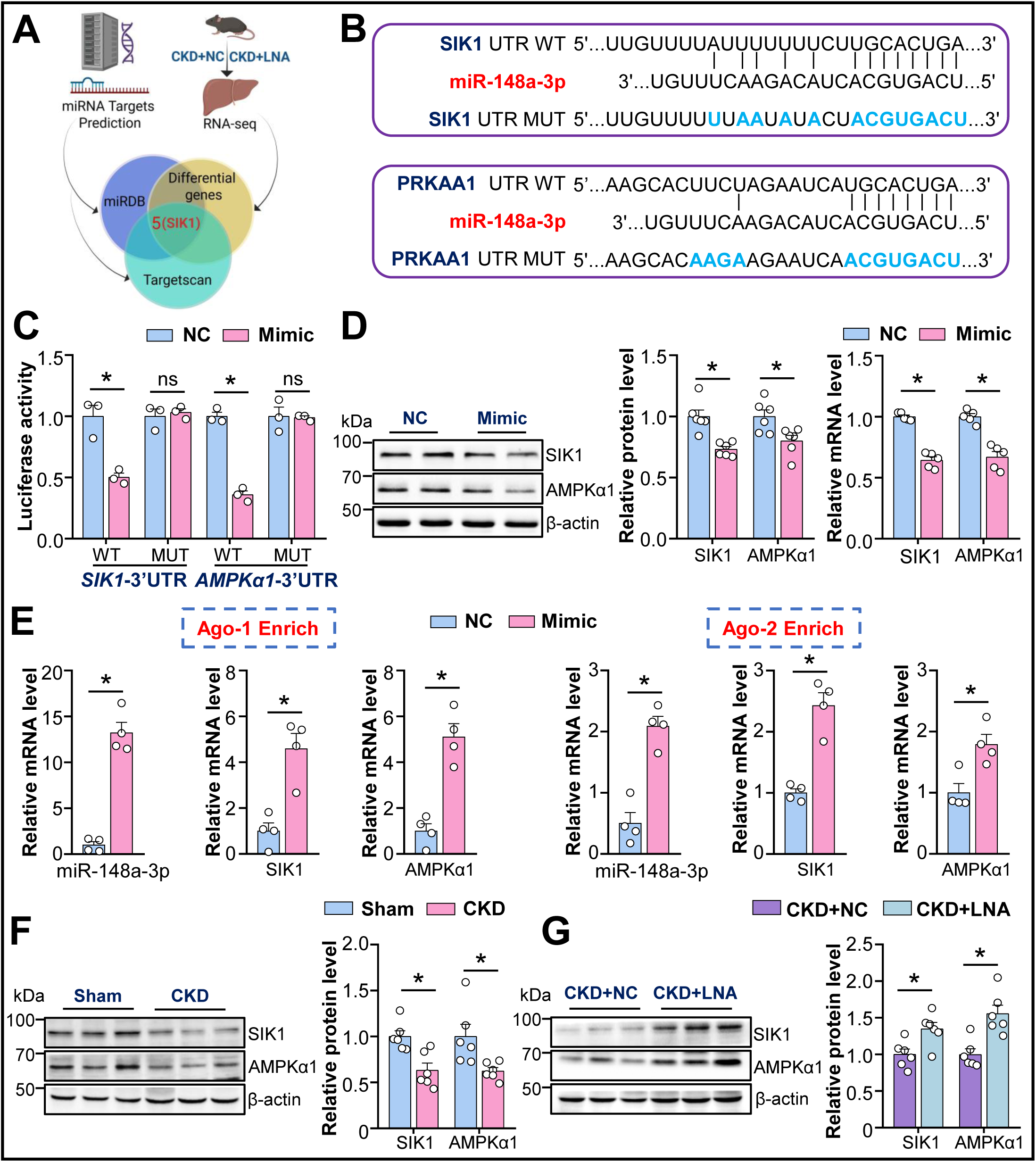
MiR-148a-3p Upregulation of SREBP2 via Targeting SIK1 and AMPKα1. **(A)** Venn diagram showing *SIK1* as a target of miR-148a-3p (miRDB - MicroRNA Target Prediction Database, purple; https://www.targetscan.org/vert_80/, green) in liver of ApoE^-/-^/nephrectomy mice administered LNA-miR-148a-3p (RNA-seq, yellow). **(B, C)** HEK293 cells were co-transfected with miR-148a-3p mimic and a luciferase reporter fused with the 3′-UTR of *SIK1* or *AMPKα1* (Luc-*SIK1*-3′-UTR wild type [WT]; *AMPKα1*-3′-UTR WT) or mutant reporters with mutated miR-148a-3p binding site (Luc-*SIK1*-3′-UTR MUT; *AMPKα1*-3′-UTR MUT). Firefly and Renilla luciferase activity were measured (n=3). **(D)** HepG2 cells were transfected with miR-148a-3p or vehicle. Protein and mRNA levels of SIK1 and AMPKα1 were assessed (n=6 for protein levels; n=5 for mRNA levels). **(E)** Immunoprecipitation of Argonaute-1 (Ago-1) or Ago-2; qPCR quantification of miRISC-associated miR-148a-3p, *Sik1*, and *Ampkα1* mRNA levels (n=4). **(F, G)** Western blot analysis of SIK1 and AMPKα1 in liver of ApoE^-/-^/nephrectomy mice and sham-operated controls; ApoE^-/-^/nephrectomy mice were administered LNA-miR-148a-3p or vehicle (n=6). Normally distributed data were analyzed by 2-tailed Student *t* test (C, D, SIK1 in F, G) or 2-tailed Student t test with Welch correction (AMPKα1 in F). Non-normally distributed data in (E) were analyzed by Mann-Whitney *U* test. \**P* < 0.05, ns=not significant.

### MiR-148a-3p/SIK1/AMPKα1 Elevates LDL-C Level via SREBP2

Because SIK1 and AMPKα1 can phosphorylate and thereby suppress SREBP1/2,^21,22^ we focused on miR-148a-3p–SIK1–SREBP2 and miR-148a-3p–AMPKα1–SREBP2 axes owing to their possible link between CKD and hyperlipidemia. To test this, we first overexpressed miR-148a-3p in HepG2 cells: levels of mature-SREBP2 and PCSK9 were increased, and that of LDLR was decreased (Figure 5A). Immunostaining confirmed mature-SREBP2 nuclear translocation driven by miR-148a-3p overexpression (Figure 5B). Next, we incubated HepG2 cells with exosomes from ASCVD/CKD or ASCVD patients and examined cellular uptake of PKH26-labled exosomes by immunofluorescent microscopy (Figure 5C). Like miR-148a-3p overexpression, exosomes from ASCVD/CKD patients significantly decreased LDLR level, possibly due to the reduced levels of SIK1 and AMPKα1, thus augmenting mature-SREBP2 and PCSK9 levels (Figure 5D). These changes were reversed in HepG2 cells transfected with LNA-miR-148a-3p (Figure 5E). In agreement with the decreased level of LDLR, the binding of fluorescence-labeled LDL was decreased in HepG2 cells overexpressing miR-148a-3p, and *SIK1* or *AMPKα1* overexpression reversed this outcome (Figure 5F). In agreement, exosomes from ASCVD/CKD patients decreased LDL binding to HepG2 cells, which was reversed by treatment with LNA-miR-148a-3p or PCSK9 monoclonal antibody (alirocumab) (Figure 5G). Together, experiments involving gain- and loss-of-function of miR-148-3p, SIK1, and AMPKα1 established the regulation of hepatic LDL metabolism by CKD-derived miR-148-3p.

**Figure 5.**
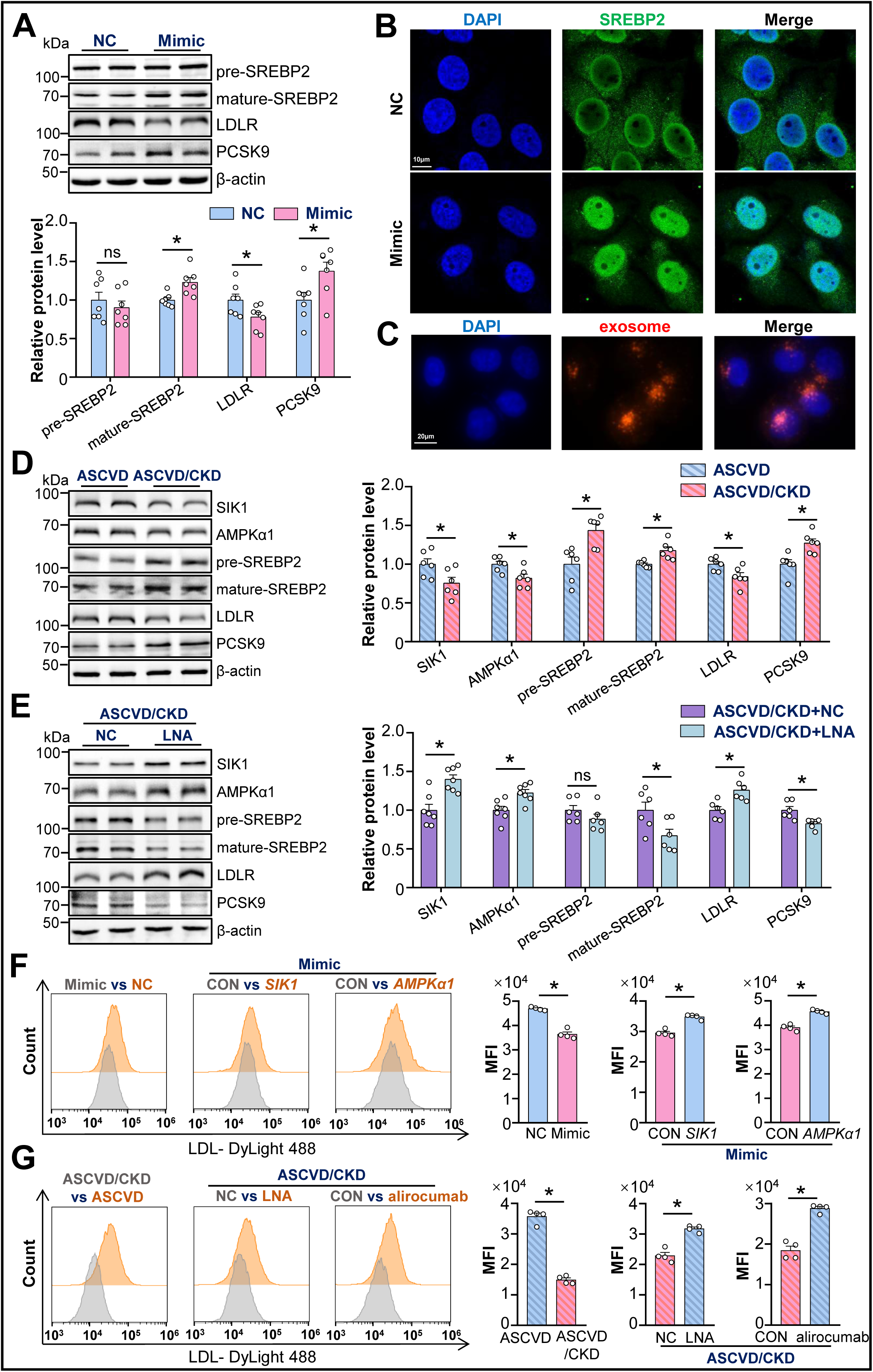
MiR-148a-3p/SIK1/AMPKα1-SREBP2 Axis Elevates LDL-C Level. **(A,B)** HepG2 cells were transfected with miR-148a-3p or vehicle for 48 h. **(A)** Protein levels of SREBP2, LDLR, and PCSK9 were assessed by western blot analysis. (n=6) **(B)** Confocal immunofluorescent microscopy of SREBP2 subcellular distribution (green); nuclei were counterstained with DAPI (blue). Scale bar: 10 μm. **(C)** Plasma exosomes from ASCVD patients with or without CKD were labeled with PKH26 fluorescence dye, incubated with HepG2 cells for 24 h, then fixed for fluorescence microscopy. Scale bar: 20 μm. (**D, E**) HepG2 cells were incubated with plasma exosomes from ASCVD patients with or without CKD. Exosome-incubated HepG2 cells were then transfected with LNA-miR-148-3p or control miRNA for 48 h. Protein levels of SIK1, AMPKα1, SREBP2, LDLR and PCSK9 were assessed by western blot analysis (n=6-7). **(F,G) (F)** HepG2 cells were transfected with miR-148a-3p, control miRNA, and expression plasmid encoding *SIK1* or *AMPKα1* as indicated (n=4). **(G)** HepG2 cells were incubated with plasma exosomes from ASCVD patients with or without CKD (left panel); In separate sets of experiments, HepG2 cells were incubated with plasma exosomes from ASCVD/CKD patients, then transfected with LNA-miR-148-3p or control miRNA (middle panel); and incubated with or without PCSK9 monoclonal antibody (alirocumab) (right panel) (n=4). At 48 h after transfection, LDL-DyLight 488 was added to all cells in (F, G) and binding to HepG2 cells was assessed by flow cytometry. Normally distributed data were analyzed by 2-tailed Student *t* test (A; SIK1, AMPKα1, LDLR and PCSK9 in D; SIK1, AMPKα1, pre-SREBP2, mature-SREBP2 and LDLR in E) or 2-tailed Student t test with Welch correction (mature-SREBP2 in D). Non-normally distributed data in (pre-SREBP2 in D; PCSK9 in E; F and G) were analyzed by Mann-Whitney *U* test. \**P* < 0.05, ns=not significant.

### SIK1/AMPKα1 Reduces Atherosclerosis in ApoE^-/-^/Nephrectomy Mice

With the established miR-148a-3p/SIK1/SREBP2 axis involved in ASCVD/CKD, we then compared the effect of hepatic overexpression of *SIK1* transcripts with or without a 3′-UTR on LDL metabolism and atherosclerosis. ApoE^-/-^/nephrectomy mice fed an HFD were administered AAV-null, AAV8-*Sik1*-3′-UTR, or AAV8-*Sik1* (Figure 6A). Among the three groups of mice, those with AAV8-*Sik1* administration had the highest hepatic level of Sik1 and Ldlr but the lowest levels of mature-Srebp2 and Pcsk9 (Figure S3C). With a moderate level of Pcsk9 in circulation (Figure 6E), these mice had the lowest levels of TG, TC, and LDL-C, consistent with reduced atherosclerosis (Figure 6B and 6C). Additionally, hepatic lipid deposition was lessened by AAV8-*SIK1* versus AAV8-*SIK1*-3′-UTR or AAV-null administration (Figure 6D), thereby suggesting the SIK1 suppression of SREBP1 as well. Given that miR-148-3p also targeted *AMPKα1* 3′-UTR, LDL-C level and atherosclerosis were mitigated in parallel experiments involving ApoE^-/-^/nephrectomy mice administered AAV8-*Ampkα1* (Figure 6F-6J). This SIK1- and AMPKα1-mitigated LDL-C level and atherosclerosis was also found in female ApoE^-/-^/nephrectomy mice (Figure S4A-D). Together, animal experiments in Figure 6 and Figure S4 recapitulated that miR-148-3p–targeted *SIK1*/*AMPKα1* might lead to hyperlipidemia and atherosclerosis in ASCVD/CKD patients.

**Figure 6.**
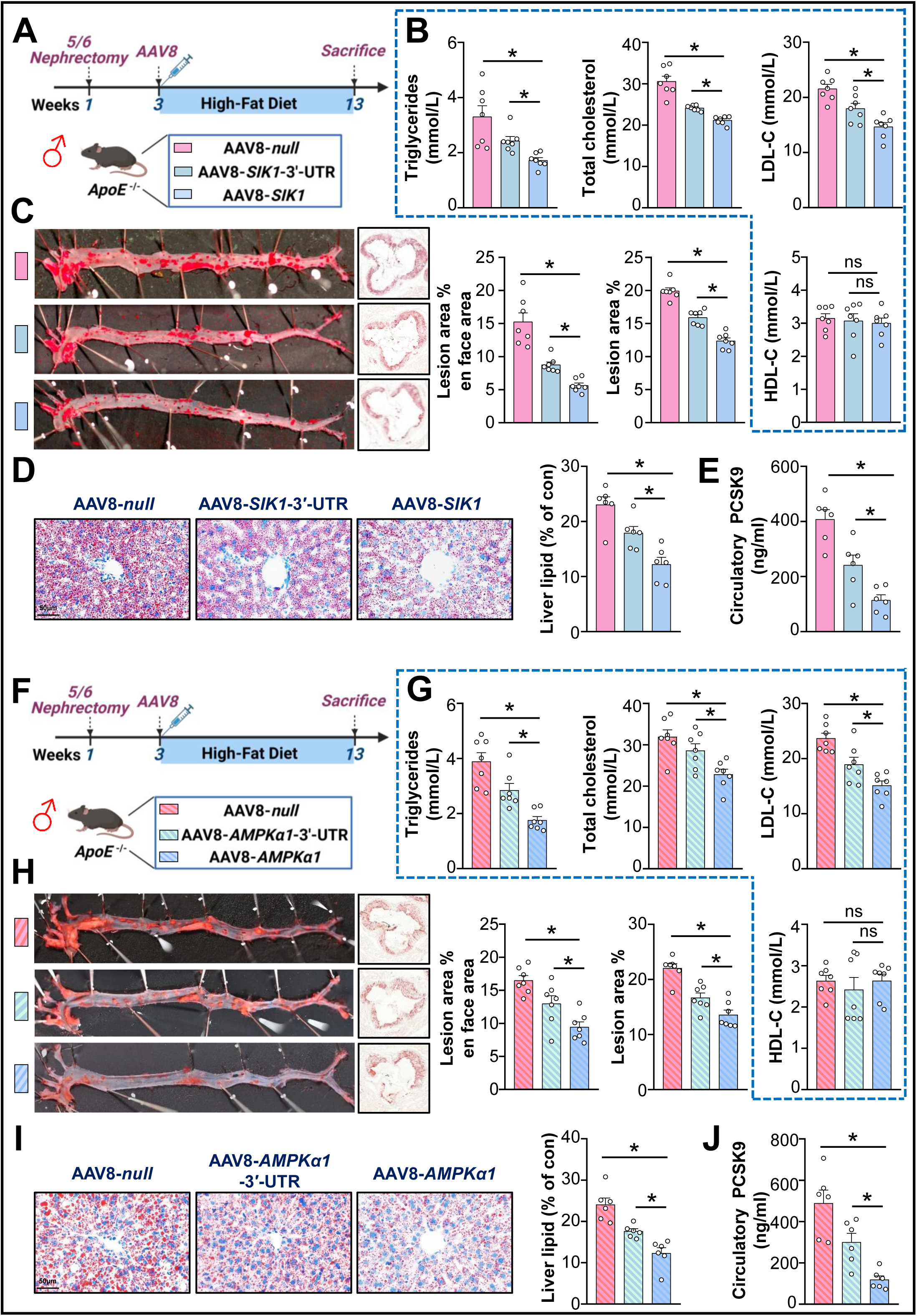
Hepatic Overexpression of SIK1/AMPKα1 Reduces Atherosclerosis in ApoE^-/-^/Nephrectomy Mice. **(A,F)** ApoE^-/-^/nephrectomy male mice were administered a single dose of AAV-null, AAV8-*Sik1*/*Ampkα1*-3′-UTR, or AAV8-*Sik1*/*Ampkα1* (2 × 10^11^ vg/mouse) by tail-vein injection and fed an HFD for 10 weeks. **(B, G)** Serum levels of TC, TG, LDL-C, and HDL-C in male mice. **(C, H)** Oil-red O *en face* staining of atherosclerosis in mouse aorta. **(D, I)** Oil-red O staining of mouse livers. **(E, J)** Plasma level of PCSK9 measured by ELISA. Data are mean±SEM from 6-7 mice per group. Data were analyzed by one-way ANOVA (LDL-C and HDL-C in B; aortic root in C; D, E, H, I and J; TG, TC and LDL-C in G), Brown-Forsythe and Welch ANOVA test (TG and TC in B; aorta in C), and Kruskal-Wallis test (HDL-C in G) among multiple groups. \**P* < 0.05, ns=not significant.

With animal experiments in Figure 6 confirming that miR-148-3p targets *SIK1*/ *AMPKα1*, leading to hyperlipidemia and atherosclerosis in ASCVD/CKD patients, one might wonder the tissue source(s) of the pathogenic miR-148-3p. To explore this, we compared miR-148a-3p level in liver, kidney, white adipose tissue (WAT), and brown adipose tissue (BAT) of ApoE^-/-^/nephrectomy mice and ApoE^-/-^/sham mice. Level of miR-148a-3p was most elevated in kidney after 5/6 nephrectomy (Figure S5A). More specifically, miR-148a-3p level was mainly increased in the tubular region of the kidney (Figure S5B). These findings suggest that damaged renal tubular cells could be a tissue source of miR-148a-3p. Packaged in exosomes, this CKD-associated miRNA can have an endocrine-like effect on hepatic LDL metabolism and atherogenesis in the arterial tree.

## Discussion

In this study, we demonstrated the role of the miR-148a-3p/SIK1/AMPKα1 axis in linking ASCVD and CKD. The new findings are that (1) miR-148a-3p level was markedly elevated in plasma exosomes from ASCVD/CKD patients and ApoE^-/-^/nephrectomy mice; (2) LNA-miR-148a-3p ameliorated hyperlipidemia and atherosclerosis in ApoE^-/-^/nephrectomy mice; (3) miR-148a-3p targeted the 3′-UTR of *SIK1* and *AMPKα1* mRNA, thus increasing SREBP2 transactivation of PCSK9 to downregulate LDLR in hepatocytes; and (4) AAV8-mediated *SIK1* and *AMPKα1* overexpression alleviated hyperlipidemia and atherosclerosis in ApoE^-/-^/nephrectomy mice. The key mechanism relies on miR-148a-3p directly targeting the 3′-UTR of *SIK1* and *AMPKα1* mRNA, which ultimately increased LDL-C level and atherosclerosis (Figure 7). Our findings provide novel insights into hyperlipidemia in CKM syndrome, revealing that miR-148a-3p is detrimental to the kidney-liver-artery axis.

**Figure 7.**
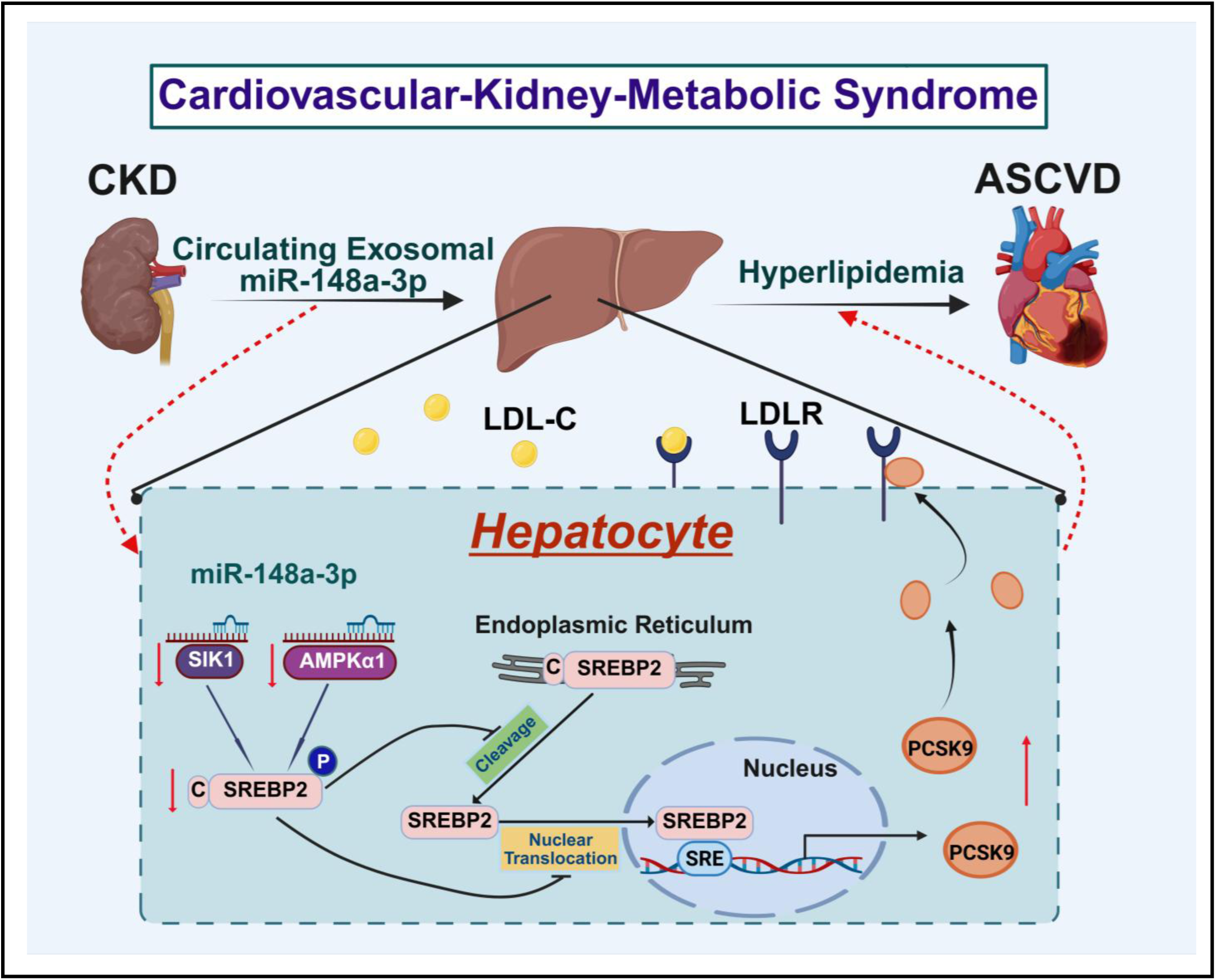
Schematic Illustration of miR-148a-3p/SIK1/AMPKα1-SREBP2 Axis Exacerbating Hyperlipidemia and Atherosclerosis. In ASCVD/CKD patients and ApoE^-/-^/nephrectomy mice, kidney-secreted miR-148a-3p targets hepatic SIK1 and AMPKα1 mRNA to activate SREBP2 and its effector PCSK9, which leads to elevated circulatory level of LDL-C and exacerbated atherosclerosis. LNA-miR-148a-3p or AAV8-*SIK1*/*AMPKα1* administration can alleviate the ASCVD/CKD-associated hyperlipidemia and atherosclerosis.

As an independent risk factor for ASCVD, CKD progression accelerates atherosclerotic plaque formation via inflammation, oxidative stress, and various metabolic abnormalities at the system level.^23,24^ Lipid metabolic disorders are common in CKD and include altered lipoprotein structure and impaired clearance, increased lipid synthesis, and compromised reverse cholesterol transport.^5,25^ Importantly, KDIGO guidelines recommend the mitigation of LDL-C levels to lower the risk of cardiovascular events in CKD patients.^25^ Clinical administration of PCSK9 mAb is effective in lowering LDL-C level in ASCVD/CKD patients.^26 27^ Emerging evidence shows that exosomal miRNAs are biomarkers and therapeutic targets for atherosclerosis.^28,29^ Others have shown that endothelial-derived miR-92a and miR-489-3p may accelerate CKD-related atherosclerosis by provoking endothelial dysfunction.^10,30^ Here, we found elevated miR-148a-3p level in circulating exosomes in ASCVD/CKD patients and ApoE^-/-^/nephrectomy mice. Moreover, the increased miR-148a-3p level was positively correlated with TC and LDL-C levels. Preclinical experiments demonstrated that miR-148a-3p antagonism significantly decreased the circulating levels of LDL-C and PCSK9, thereby alleviating hyperlipidemia and atherosclerosis in ApoE^-/-^/nephrectomy mice. Thus, our work provides mechanistic insights into the elevated PCSK9 level in ASCVD/CKD patients and also suggests that miR-148a-3p can be a new drug target, in addition to PCSK9 mAb and inclisiran, used for lipid-lowering.^25,31^

A previous study showed that targets of miR-148a-3p, including LDLR and ABCA1, might affect lipid metabolism but did not show a lipid-lowering effect of LNA-miR-148a-3p in ApoE^-/-^ mice.^19^ Moreover, the authors’ subsequent *in vivo* studies revealed that miR-148a antagonism likely reduces the macrophage-dependent mechanism rather than LDL-C metabolism.^15^ Our results suggested that LNA-miR-148a-3p mitigated atherosclerosis in ApoE^-/-^/nephrectomy mice via lipid-lowering effects. As compared with increased miR-148a-3p levels caused by the HFD alone, nephrectomy-increased nephrogenic miR-148a-3p level was delivered to the liver by circulatory exosomes, which exacerbated LDL-C. This finding differs from the mechanism proposed by the previous study. Mechanically, our study suggested that the CKD-derived miR-148a-3p worsened hyperlipidemia and atherosclerosis by targeting AMPK family kinases (SIK1/AMPKα1) that are the principal regulators of SREBP2-mediated lipid and cholesterol metabolism. *In vitro*, overexpression of *SIK1*/*AMPKα1* or PCSK9 mAb administration reversed the miR-148a-3p–decreased LDL binding, so miR-148a-3p regulates LDL-C and ASCVD in a multi-factorial manner. Thus, our study suggests that pharmacologic interventions for hyperlipidemia and ASCVD/CKD can be achieved by targeting the miR-148a/SIK1/AMPK-SREBP2 axis. These interventions include statins, PCSK9 mAb, metformin (for AMPK/SIK1 activation), and LNA-miR-148a, alone or combined.

Functioning as an energy sensor, AMPK regulates hepatic cholesterol metabolism by phosphorylating and thereby inhibiting SREBP2 transactivation of downstream target genes such as HMG-CoA reductase.^22,32^ SIK1, as a member of the AMPK-related kinase family, can modulate hepatic triglyceride synthesis via phosphorylation of SREBP1-c.^21,33^ Although pivotal for cholesterol and fatty acid metabolism, AMPK and SIK1 play important roles in vascular health via phosphorylation of substrates such as eNOS, ACE2, and KLF2.^34-37^ Although the 3-kb upstream regions of human *PRKAA1* and *SIK1* genes and their 500-bp 3′-UTR are 46% and 46% homologous, their miR-148a-3p seed sequences are identical. Given the downregulated AMPK and SIK1 in the vasculature of ASCVD/CKD, common regulatory mechanisms, including miR-148a-3p 3′-UTR targeting, post-translational modifications, and transcriptional suppression, would collectively contribute to this downregulation. With downregulated AMPK and SIK1, the hepatic level of LDLR is decreased by SREBP2-transactivated PCSK9. With AMPK and SIK1 positively but PCSK9 negatively regulating vascular health, a therapeutic strategy involving miR-148a-3p antagonism would downregulate PCSK9 in the circulation and liver and also upregulate beneficial genes in the vasculature. This notion is supported by SIK1 or AMPKα1 overexpression decreasing plasma levels of PCSK9 and LDL-C as well as atherosclerosis in our ApoE^-/-^/nephrectomy mice (Figure 6).

In addition to alterations in miR-148a-3p level, CKD induces alterations in miR-16-5p, miR-17-5p, miR-20a-5p, and miR-106b-5p levels, which target the VEGFA-VEGFR2 signaling pathway to enhance vascular calcification.^9^ Our results in Figure S5 show that CKD robustly increased miR-148a-3p level in the kidney, in line with a previous study of rats undergoing unilateral ureteral ligation.^38^ On further analysis, miR-148a-3p was significantly enriched in the mouse tubular regions after 5/6 nephrectomy (Figure S5B). Also, uremic toxins increased miR-148a-3p level in cultured HK2 cells (Figure S5C). Thus, CKD led to an elevated level of miR-148a-3p in renal tubular epithelial cells. The renal-originated miR-148a-3p then entered circulation via exosomes, ultimately targeting AMPK and SIK1 in the liver and vasculature and contributing to lipid metabolism disorders and vascular impairments. Intriguingly, the circulatory level of miR-148a-3p in ASCVD/CKD patients was inversely correlated with eGFR and positively with TC and LDL-C levels (Figure 1). The inverse correlation of miR-148a-3p level and eGFR also occurred in urine specimens from ASCVD patients with CKD (Figure 1). Thus, miR-148a-3p could serve as a biomarker for cardiovascular risk for ASCVD/CKD.

In summary, this study showed that CKD-induced miR-148a-3p is pathogenic for hyperlipidemic disorders and ASCVD. This endocrine-like effect between the kidney, liver, and arterial wall may be a sequela of CKM, which points to miR-148a-3p as a therapeutic target of ASCVD/CKD.

## Non-standard Abbreviations and Acronyms

3’-UTR: 3’ untranslated region
AAV8: adeno-associated virus serotype 8
AMPKα1: AMP-activated protein kinase alpha 1 subunit
ASCVD: atherosclerotic cardiovascular disease
CKD: chronic kidney disease
CKM: cardiovascular-kidney-metabolic
eGFR: estimated glomerular filtration rate
FISH: fluorescence in situ hybridization
HDL-C: high-density lipoprotein cholesterol
HFD: high fat diet
LDL-C: low-density lipoprotein cholesterol
LDLR: low-density lipoprotein receptor
miRISCs: miRNA-induced silencing complexes NTA nanoparticle tracking analysis
PCSK9: proprotein convertase subtilisin/kexin type 9
SIK1: salt-inducible kinase 1
SREBP-1c: sterol regulatory element binding protein-1c
SREBP2: sterol regulatory element binding protein 2
TC: total cholesterol
TEM: transmission electron microscopy
TG: triglycerides

## Acknowledgments

We thank Drs. Junhui Liu and Liyi Xie at First Affiliated Hospital of Xi’an Jiaotong University, and Min Chen at Peking University First Hospital for their technical assistance and consultation. The authors acknowledge the use of BioRender for creating the schematic.

## Author Contributions

Y.H., T.L., E.H., P.Z., J.Y.-J.S., Z.-Y.Y., and J.Z. conceived of the study and designed the experiments. Y.H., T.L., E.H., P.Z., X.F., H.F., Z.W., Y.L., C.W., W.Y., L.W., P.L. and B.D. performed the experiments. Y.H. and Y.L. conducted in silico analysis. Y.H., T.L., E.H., P.Z., X.F., Y W, W.L., J.F., X.B., H.W., J.L., J.Y.-J.S., Z.-Y.Y., and J.Z. analyzed the data and provided the discussion. Y.H., T.L., E.H., P.Z., J.Y.-J.S., Z.-Y.Y., and J.Z. wrote the manuscript. All authors reviewed the paper and approved the final draft.

